# Exploring the Changing Landscape of Cell-to-Cell Variation After CTCF Knockdown via Single Cell RNA-seq

**DOI:** 10.1101/862847

**Authors:** Wei Wang, Gang Ren, Ni Hong, Wenfei Jin

## Abstract

**Background:** CCCTC-Binding Factor (CTCF), also known as 11-zinc finger protein, participates in many cellular processes, including insulator activity, transcriptional regulation and organization of chromatin architecture. Based on single cell flow cytometry and single cell RNA-FISH analyses, our previous study showed that deletion of CTCF binding site led to a significantly increase of cellular variation of its target gene. However, the effect of CTCF on genome-wide landscape of cell-to-cell variation is unclear.

**Results:** We knocked down CTCF in EL4 cells using shRNA, and conducted single cell RNA-seq on both wild type (WT) cells and CTCF-Knockdown (CTCF-KD) cells using Fluidigm C1 system. Principal component analysis of single cell RNA-seq data showed that WT and CTCF-KD cells concentrated in two different clusters on PC1, indicating gene expression profiles of WT and CTCF-KD cells were systematically different. Interestingly, GO terms including regulation of transcription, DNA binding, Zinc finger and transcription factor binding were significantly enriched in CTCF-KD-specific highly variable genes, indicating tissue-specific genes such as transcription factors were highly sensitive to CTCF level. The dysregulation of transcription factors potentially explain why knockdown of CTCF lead to systematic change of gene expression. In contrast, housekeeping genes such as rRNA processing, DNA repair and tRNA processing were significantly enriched in WT-specific highly variable genes, potentially due to a higher cellular variation of cell activity in WT cells compared to CTCF-KD cells. We further found cellular variation-increased genes were significantly enriched in down-regulated genes, indicating CTCF knockdown simultaneously reduced the expression levels and increased the expression noise of its regulated genes.

**Conclusions:** To our knowledge, this is the first attempt to explore genome-wide landscape of cellular variation after CTCF knockdown. Our study not only advances our understanding of CTCF function in maintaining gene expression and reducing expression noise, but also provides a framework for examining gene function.

## Background

CCCTC-binding factor (CTCF) is an 11-zinc finger protein that directionally binds to a well-defined DNA motif [1, 2]. Although CTCF was initially reported as a transcription factor [3, 4], subsequent studies found it served as an insulator [5–7]. Nowadays, CTCF has been reported being involved in multiple cellular processes, such as transcriptional regulation, insulator activity, epigenetic regulation, organization of chromatin architecture and X chromosome inactivation [1, 2, 8–14]. CTCF activates and silences gene expression by preventing the spread of heterochromatin and blocking of unrelated enhancer-promoter interactions [1, 15]. Interestingly, mammalian genome is organized into thousands of highly self-interacting topologically associated domains (TADs) with CTCF demarcating individual TAD boundary [16]. Analyses based on high-resolution interaction matrix further identified ~10,000 chromatin loops ranging ~185Kb in human genome, anchoring by convergent CTCF-binding motif-pair at TAD boundaries [13]. In addition, chromatin loops mediated by CTCF and cohesin can tether distal enhancers to gene promoters and regulate its target gene expression [13, 14, 17, 18]. Further studies showed CTCF-mediated DNA loop could determine the chromatin architecture, with anchor-genes almost exclusively being housekeeping genes, while loop-genes being tissue-specific genes [14, 19]. For instance, inversion of a CTCF-binding site reconfigured the topology of chromatin loops and activated gene expression by creating a new chromatin loop [20, 21].

Recent studies showed that disruption of CTCF binding in mammalian cells resulted in loss of TADs [18], genomic instability [22], developmental failure [23, 24] and other malfunctions. Our recent study demonstrated that CTCF played an important role in stabilizing enhancer-promoter interaction and reducing the gene expression noises in mammalian cells [17]. In particular, we found that CTCF-KD or deletion of CTCF binding sites led to increased variation of cellular expression of GATA3, CD28, CD90 and CD5 [17]. However, the genome-wide change of cell-to-cell variation after CTCF-KD remains unknown. In this study, we conducted single cell RNA-seq on both WT and CTCF-KD cells to investigate the changing landscape of cell-to-cell variation at a genome-wide scale. Interestingly, GO terms including regulation of transcription, DNA binding and Zinc finger were significantly enriched in CTCF-KD specific highly variable genes. We also found that cellular variation-increased genes were significantly enriched in down-regulated genes, indicating knockdown of CTCF simultaneously reduced the expression level and increased the expression noises of its regulated genes.

## Results

### Efficient CTCF knockdown and single cell RNA-seq

We knocked down CTCF in EL4 cells by short hairpin RNA (shRNA). Western blotting showed a dramatic decrease of the CTCF protein level in shCTCF #1 and shCTCF #2 compared to shRNA luciferase controls (shLuc) (Fig. 1A). Quantitative polymerase chain reaction (qPCR) revealed the mRNA levels in shCTCF #1 and shCTCF #2 had been reduced to 38% and 40% of that in shLuc, respectively (Fig. 1B). These results confirmed the efficient knockdown of CTCF expression in shCTCF#1 and shCTCF#2, with a significant reduction consistently at both RNA and protein level.

**Fig 1.**
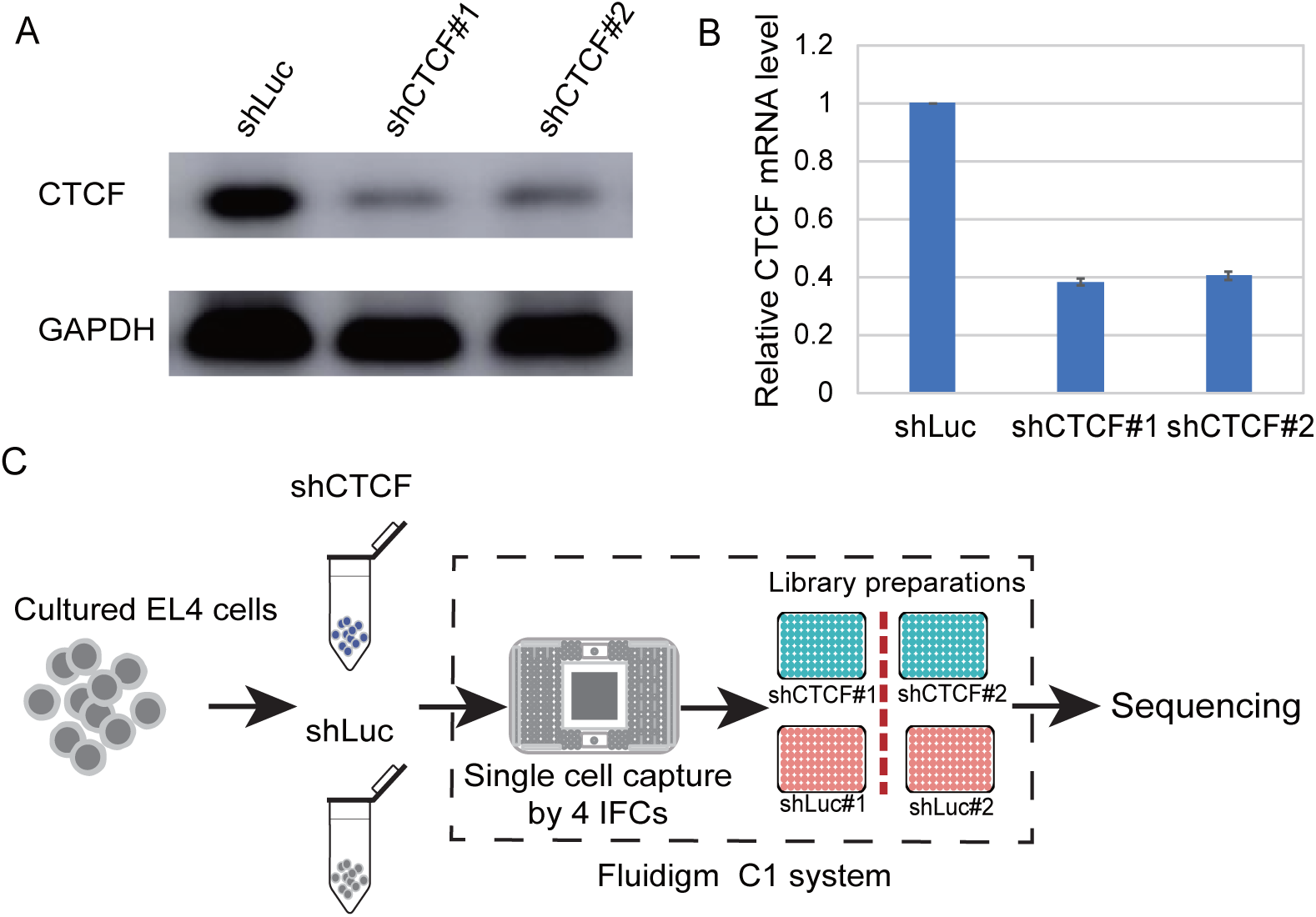
Knockdown of CTCF and schema of single cell sequencing. **A.** Western blot analysis of CTCF in luciferase control (shLuc) and CTCF-KD cells (shCTCF#1 and shCTCF#2). EL4 cells were infected with retroviral particles encoding GFP and an shRNA targeting CTCF or a control sequence for 5 days. **B.** Real-time quantitative PCR (RT-qPCR) analysis of CTCF expression in luciferase control (shLuc) and knockdown (shCTCF#1 and shCTCF#2) cells. The expression level of CTCF was normalized to GAPDH. **C.** Schema of single cell RNA sequencing using Fluidigm C1 system.

In order to investigate the changing landscape of cell-to-cell variation after CTCF knockdown, we successfully conducted single cell RNA-seq for shLuc#1, shLuc #2, shCTCF #1 and shCTCF#2 using 4 integrated fluidics circuits (IFCs) (Fig. 1C). We noticed the gene expression level of pooled shLuc single cells were highly correlated with that of bulk data from our previous study [17] (r^2^=0.86; Fig. S1A). The gene expression of pooled single cells from shLuc #1 was also highly correlated with that of shLuc #2 (r^2^=0.87; Fig. S1B). In addition, the gene expression of pooled single cells repeats and bulk cell repeats in CTCF-KD cells were highly correlated (Fig. S1C-S1D).

### Systematic differences between WT cells and CTCF-KD cells

A total of 95 cells, including 24 cells from shLuc #1, 24 cells from shCTCF #1, 23 cells from shLuc #2 and 24 cells from shCTCF #2, were kept for further analyses after quality control. We conducted principal component analysis (PCA) on 11,361 genes shared by those 95 cells (Fig. 2A). The coordination of cells from experiment1 and experiment2 on PCA projection is not significantly different (P=0.8; student’s t-test), indicating no obvious batch effect between experiment1 and experiment2. Further analysis showed that WT cells and CTCF-KD cells were distinguishable on PCA projection and concentrated in two different clusters on PC1 (Fig. 2B, 2C), implying the gene expression profiles of CTCF-KD and WT cells were systematically different. We also noticed a correlation between CTCF expression level and its coordination on the PCA projection (using the first 10 PCs), among which CTCF expression level and PC2 exhibited the highest correlation (Fig. S2A) (r^2^=0.18, P = 0.22×10^−6^).

**Fig 2.**
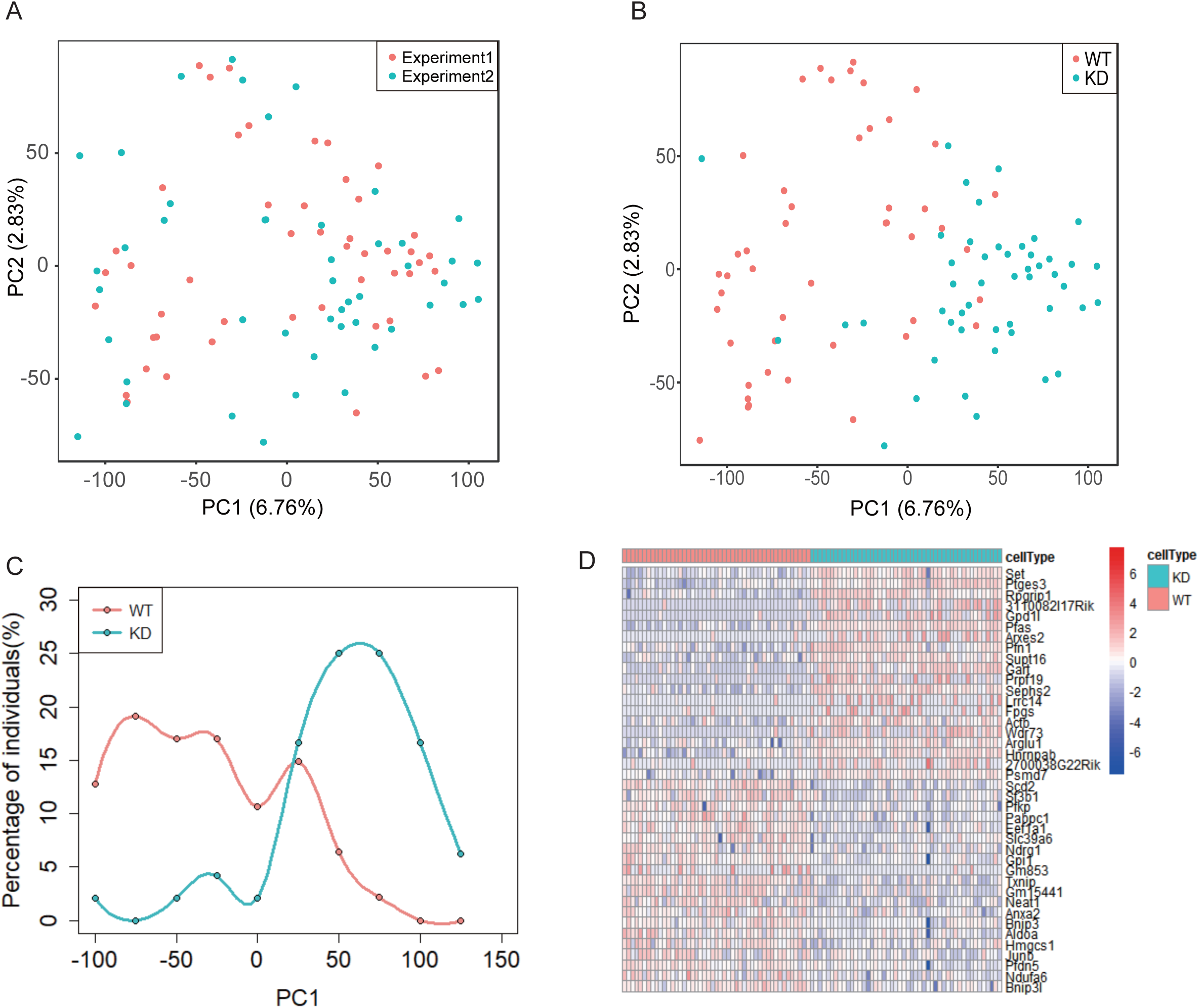
There is systematic difference between CTCF-KD and WT cells. **A.** No significant batch effect among the experimental repeats based on PCA analysis. **B.** CTCF-KD cells were largely distinguishable from WT cells on PCA projection. **C.** Distribution of individual WT cells and individual CTCF-KD cells on PC1. **D.** Heatmap of differentially expressed genes (TOP 20) between WT cells and CTCF-KD cells.

We further calculated the differential gene expression between WT and CTCF-KD cells using edgeR [25]. We identified 195 up-regulated and 107 down-regulated genes in CTCF-KD cells compared to WT cells (Fig. S2B). Heatmap of the most differentially expressed genes between WT cells and CTCF-KD cells exhibited a cellular heterogeneity within the same cell population (Fig. 2D). The most enriched gene categories in down-regulated genes include glycolytic processing, Prolyl 4-hydroxylase α subunit, iron-dependent dioxygenase and carbon metabolism (Fig. S2D). Whereas the most enriched gene categories in up-regulated genes include RNA binding, ribosome biogenesis, WD40 repeat domain and RNA processing (Fig. S2C), consistent to our recent study based on bulk data in some way [17].

### CTCF knockdown changed the landscape of cell-to-cell variation

In order to distinguish true signals of cellular variation from technical noise, we calculated the expression noise of each gene (σ^2^/μ^2^) [26, 27]. The expression noises exhibited two distinct scaling properties: negative association with expression at low expression levels and no association at high expression levels (log_2_TPM>1) (Fig. 3A). We filtered out low expressed genes (log_2_TPM≤1) to reduce the impact of technological noise, resulting in 7,843 genes for further analysis (Fig. 3A). Coefficient of variation (CV) was calculated to measure the cell-to-cell variation of each gene across the cell populations. The distribution of changes of cell-to-cell variation pre- and post-CTCF knockdown followed normal distribution (Fig. S3). We identified 602 cellular variation increased genes and 890 cellular variation decreased genes after CTCF knockdown by mean±SD (Fig. 3B).

**Fig 3.**
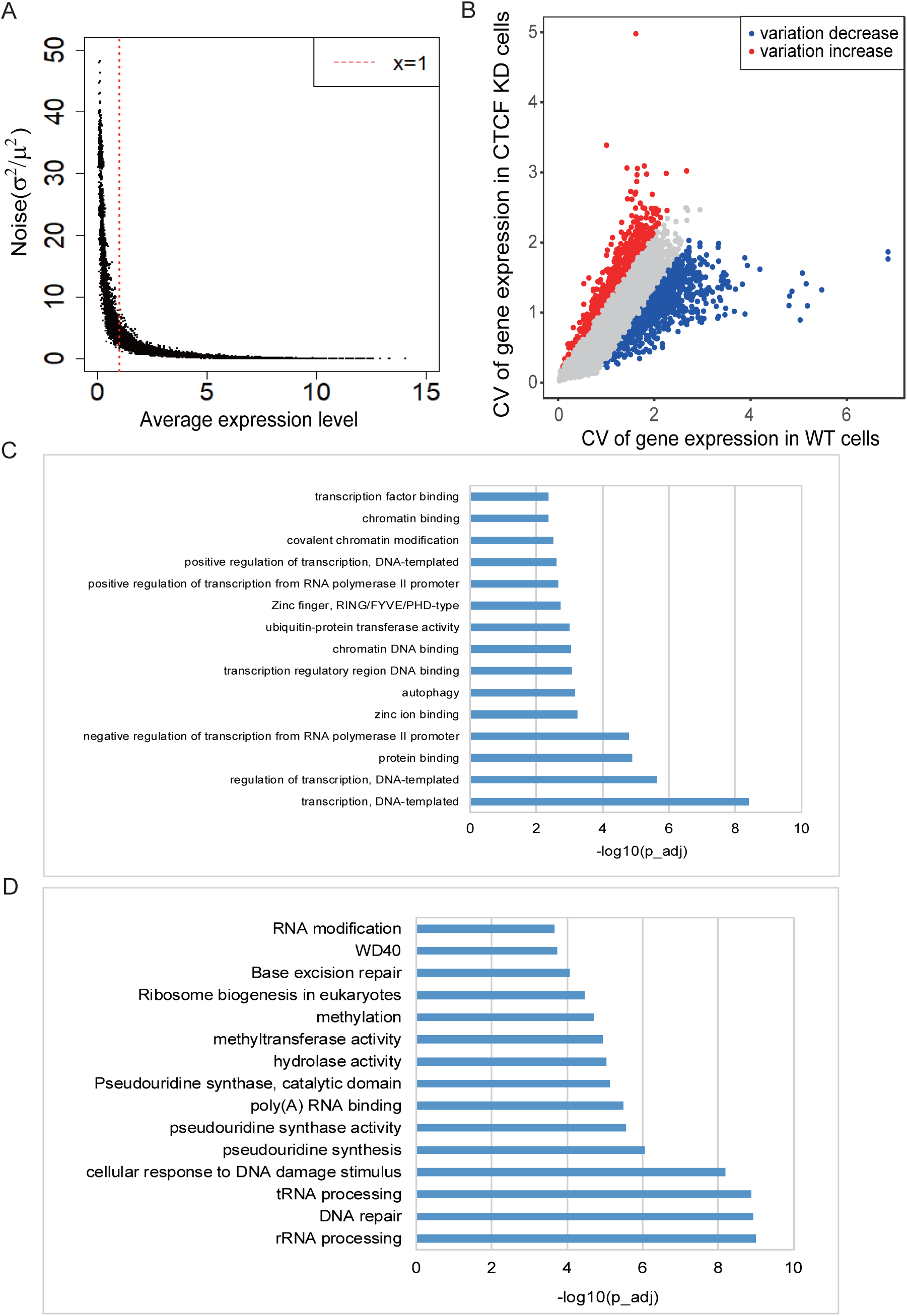
Identification and analyses of genes showing cellular variation-changed after CTCF KD. **A.** The relationship between expression level and noise level of reference genes. Genes with low cellular variation was used for further analyses. **B.** Scatter plot of the cellular variation-changed genes after CTCF KD. Blue and red indicate the variation decrease and variation increase, respectively. **C.** The top 15 gene categories enriched in variation increased gene. **D.** The top 15 gene categories enriched in variation decreased gene.

GO analyses showed that variation-increased genes were significantly enriched in GO terms such as regulation of transcription, DNA binding, zinc finger proteins, covalent chromatin modification and transcription factor binding (Fig. 3C). In fact, almost all genes in DNA binding, zinc finger proteins and transcription factor binding are transcription factors. The significant enrichment of transcription factors in CTCF-KD-specific highly variable genes potentially indicates that transcription factors are highly sensitive to CTCF level and are tend to be cellular variation increased genes. The dysregulation of a lot transcription factor potentially explain why knockdown of CTCF lead to systematic change of gene expression. In contrast, variation-decreased genes were significantly enriched in housekeeping genes related GO terms such as rRNA processing, DNA repair, tRNA processing, and RNA modification (Fig. 3D). The enrichment of housekeeping genes in WT-specific highly variable genes potentially indicates a higher cellular variation of cell activity in WT cells compared to CTCF-KD cells.

### CTCF Knockdown simultaneously altered expression levels and cellular variations of its regulated genes

We identified 302 expression-changed genes and 1,490 cellular variation-changed genes pre-and-post CTCF knockdown. It is interesting to examine whether those cellular variation-changed genes were enriched in expression-changed genes. Venn diagram showed that 47 genes out of total 107 down-regulated genes exhibited increased cellular variation (Fig. 4), which were significantly over-represented (P=0.29×10^−23^, χ^2^ test), indicating CTCF knockdown simultaneously reduced the expression level and increased the gene expression noise. Among those genes with decreased expression and increased cellular variation, EGR1 and JUNB played an important role in maintaining the cell type-specific gene regulation. For instance, EGR1 belongs to the EGR family of C2H2-type zinc-finger proteins, and encodes a nuclear protein that participates in transcriptional regulation.

**Fig 4.**
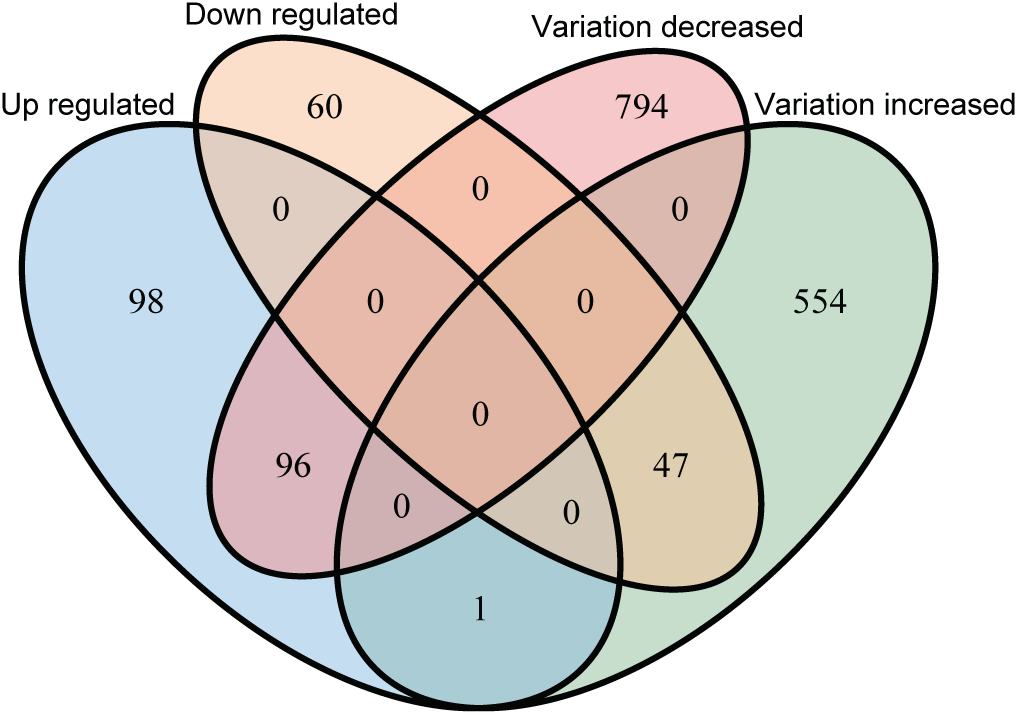
Genes showing cellular variation change are more likely to be differentially expressed genes. Genes with expression decreased and cellular variation increased were significantly over-represented (P=0.29×10^−23^, χ^2^ test). Genes with decreased cellular variation and increased expression level were significantly over-represented (P=0.48×10^−23^, χ^2^ test).

Meanwhile, there were 96 genes out of the total 195 up-regulated genes exhibiting decreased cellular variation (Fig. 4), which were also significantly over-represented (P=0.48×10^−23^, χ^2^ test). The 96 genes with decreased cellular variation and increased expression level were significantly enriched in poly(A) RNA binding, rRNA processing, WD40, purine nucleobase biosynthetic processing, rRNA methylation and RNA methyltransferase activity. It is obvious that those enriched GO terms were associated with basic cellular functions belonging to housekeeping genes. Taken together, our results clearly indicate that distortion of CTCF expression could simultaneously change the gene expression level and cell-to-cell variation of its regulated genes.

Furthermore, we identified CTCF binding sites using CTCF ChIP-seq data in WT EL4 cells from our previous study [17]. We identified each gene-associated CTCF by counting the CTCF binding sites within 20Kb of the transcriptional start site (TSS) for each gene. The numbers of gene-associated CTCF of variation-increased genes are significantly higher than that of variation-decreased genes (P= 0.0033; Wilcoxon test), and are significantly higher than that of variation-unchanged genes (P= 0.16×10^−6^; Wilcoxon test). The numbers of gene-associated CTCF of variation-decreased genes are not significantly different from that of variation-unchanged genes (P= 0.5; Wilcoxon test). These results suggest that genes regulated by multiple CTCF binding sites tend to possess higher cellular variation after CTCF knockdown.

## Discussion

CTCF plays an important role in chromatin structure organization and regulation of gene expression [13–16]. In this study, we used single cell RNA-seq to analyze genome-wide gene expression profiles of WT and CTCF-KD cells at single cell resolution. Indeed, WT cell population and CTCF-KD cell population showed distinct concentration on PC1, indicating that knockdown of CTCF resulted in a systematic impact on the genome-wide gene expression profile. These results further implied that CTCF contributed to key functions in controlling the genome-wide gene regulation. We generated the genome-wide landscape of cell-to-cell variation in both WT and CTCF-KD cells. After comparing cell-to-cell variations between WT and CTCF-KD cells, we identified those genes showing significant change of cellular variation after CTCF knockdown. Interestingly, the cellular variation-increased genes are significantly enriched in expression-decreased genes, suggesting CTCF-medicated promoter-enhancer interaction did not only play an important role in maintaining the expression of its regulated genes, but also reduced their expression noise.

In this study, we identified a lot gene with an obvious change of cellular variation after CTCF knockdown. Interestingly, the variation-increased genes were significantly enriched in GO terms such as chromatin DNA binding, zinc finger proteins and zinc ion binding, indicating the expression noise of those zinc finger proteins were strongly increased after CTCF knockdown. The increased cellular variation of zinc finger proteins potentially indicates a high sensitivity of zinc finger proteins to CTCF expression level or cellular environmental change within the cell. In fact, the majority of those zinc finger proteins were transcription factors that played an important role in the regulation of cell type-specific gene expression. Our observation that CTCF knockdown fluctuated expression of a lot transcriptional factor further explain why disruption of CTCF expression led to pronounced biological effects such as development failure [23, 24]. Taken together, our findings provide convincing evidence that CTCF serves as a key player in stabilizing the gene expression noise of zinc finger related genes.

## Conclusion

We conducted single cell RNA-seq on both wild type (WT) cells and CTCF-Knockdown (CTCF-KD) cells using Fluidigm C1 system. Principal component analysis of single cell RNA-seq data showed that WT and CTCF-KD cells concentrated in two different clusters on PC1, indicating gene expression profiles of WT and CTCF-KD cells were systematically different. Interestingly, GO terms including regulation of transcription, DNA binding, Zinc finger and transcription factor binding were significantly enriched in CTCF-KD-specific highly variable genes, indicating tissue-specific genes such as transcription factors were highly sensitive to CTCF level. The dysregulation of transcription factors potentially explain why knockdown of CTCF lead to systematic change of gene expression. In contrast, housekeeping genes such as rRNA processing, DNA repair and tRNA processing were significantly enriched in WT-specific highly variable genes, potentially due to a higher cellular variation of cell activity in WT cells compared to CTCF-KD cells. We further found cellular variation-increased genes were significantly enriched in down-regulated genes, indicating CTCF knockdown simultaneously reduced the expression levels and increased the expression noise of its regulated genes. To our knowledge, this is the first attempt to explore genome-wide landscape of cellular variation after CTCF knockdown. Our study not only advances our understanding of CTCF function in maintaining gene expression and reducing expression noise, but also provides a framework for examining gene function.

## Methods

### Cell culture

EL4 cells were cultured in DMEM (GIBCO Invitrogen) supplemented with 50 IU/mL penicillin, 50 mg/mL streptomycin (GIBCO Invitrogen) and 10% heat-inactivated calf serum (Sigma, USA). Cultures were maintained by replacement of fresh medium every 3 days, and cell density was kept between 1 × 10^5^ and 1 × 10^6^ cells/mL.

### Knockdown of CTCF by shRNA

Knockdown of CTCF was performed using Lentiviral-mediated short hairpin RNA (shRNA) in EL4 cells as described previously [17]. Briefly, 293T cells were co-transfected with an envelope plasmid (pLP/VSVG) to generate lentiviral particles. The medium containing lentiviral particles was harvested after 48 hours transfection. EL4 cells were infected with the harvested shLuc and shCTCF retroviral particles packaged in GP2-293. Some GFP+ cells were sorted out to check the knockdown efficiency using RT-qPCR and Western blotting after 5 days of infection. The cell populations displaying efficient knockdown of CTCF were used for single cell RNA-seq.

The following shRNA sequences were used for CTCF knockdown: mouse CTCF-shRNA 1: 5’-GGTGCAATTGAGAACATTATA; mouse CTCF-shRNA 2: 5’-TGGACGATACCCAGATCATAA.

### Western blot analyses

After thorough washing, the knockdown (shCTCF) cells and control (shLuc) were harvested for Western blotting analyses. Protein concentration of the cell lysates was measured using BCA kit (Pierce, Rockford, IL). Protein samples (40µg/lane) were applied to SDS-PAGE followed by Western blotting against anti-CTCF antibody (07-729, Millipore), and anti-GAPDH antibody (sc-1616, Santa Cruz Biotechnology).

### Quantitative Real-time PCR

Total RNAs from the knockdown (shCTCF#1 and shCTCF#2) and control (shLuc) cells were extracted using miRNeasy Micro Kit (QIAGEN). cDNA was synthesized by using oligo (dT)20 and SuperScript III Reverse Transcriptase (Invitrogen) according to manufacturers’ instructions. RT-qPCR samples were mixed with the following Taqman probes mixture (Applied Biosystems) and run on a LightCycler 96 (Roche): Gapdh: Mm03302249_g1; CTCF: Mm00484027_m1. Results were normalized to the mRNA level of Gapdh.

### Single cell RNA sequencing

Fluidigm C1™ Single-Cell Autoprep System (Fluidigm, South San Francisco, CA, USA) was used for single cell RNA-seq. In the initial experiment, shLuc#1 or shCTCF#1 were uploaded to a C1 integrated fluidics circuit (IFC) for cell capture, respectively. We checked the IFC to count the number of captured cells, and to distinguish between live and dead cells for later data processing. After successful completion of the second knockdown, cells from shLuc#2 and shCTCF#2 were treated similarly to the initial experiment. Single-cell RNA-seq with SMARTer protocol (Clontech, Mountain View, CA, USA) was prepared following Fluidigm manual ‘Using the C1 Single-Cell Auto Prep System to generate mRNA from Single Cells and Libraries for Sequencing’. The wells containing either zero or double cells were filtered out. We selected 24 cells with the highest quality from each IFC. The DNA materials obtained from the 96 single cells were sequenced on Illumina HiSeq 3000, as illustrated in Fig. 1C.

### Reads mapping and quality control

Quality of the reads was assessed using FASTQC. All reads were aligned to the mouse genome (Ensemble version GRCm38.89) utilizing STAR v.2.5.2 [28, 29]. Unique mapping reads were allowed (using parameter *--outFilterMultimapNmax*). The alignments were used as input in HTSEQ v.0.9.1 [30] to count the number of reads mapping to each of the 24,057 ref-seq genes in each cell. We filtered out those low-quality cells from our dataset based on a threshold for a minimum of 3000 unique genes per cell. Transcripts per million (TPM) was used to normalize the gene expression level and log2 transformed. Furthermore, genes with log2(TPM+1)>1 in less than two individual cells were filtered out, leaving a total of 95 samples and 11,361 genes for further analyses.

### Statistical analyses and gene enrichment analyses

For identification of genes with biologically significant cell-to-cell variation, we used η^2^ = σ^2^/μ^2^ (*σ* denotes standard deviation; *µ* denotes mean) to measure the noise of gene expression [26, 27, 31]. We filtered out those genes exhibiting a low expression (log_2_TPM≤1), since the expression noise is inversely proportional to the expression when gene expression level is low (log_2_TPM≤1) (Fig. 3A), leading to 7,843 genes remaining for further analyses. To examine any possible enrichment of particular gene categories and pathways in certain gene lists, GO enrichment analysis was performed using DVAID [32, 33]. GO categories with Benjamini < 0.05 were considered as significant.

In this study, we used coefficient of variation (CV) to calculate the cell-to-cell variation. The variation increased genes is calculated by

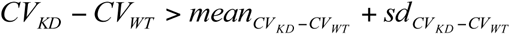

while the variation decreased genes is calculated by

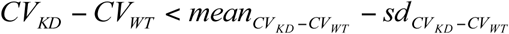

## Supporting information

Supplemental Figur1

Supplemental Figur3

## Acknowledgements

We thank members of our laboratory for their suggestions and encouragement. The computation was supported by Center for Computational Science and Engineering of Southern University of Science and Technology.

## Author’s contributions

W.J. conceived the project. R.G. performed the experiments and W.W. analyzed the data. N.H. and W.J. supervised the study. W.W., W.J. and N.H. wrote the manuscript.

## Data Availability

The single cell RNA-seq data of EL4 cells have been deposited in the Gene Expression Omnibus database with accession number GSE135769.

## Supplementary figures

**Fig S1. High reproducibility of single cell RNA-seq data.**

(A) Scatter plot showing the correlation of gene expressions between pooled single cells and bulk data in shLuc.

(B) Scatter plot showing the correlation of gene expression between shLuc #1 and shLuc#2.

(C) Scatter plot showing the correlation of gene expressions between pooled single cells and bulk data in CTCF-KD cells.

(D) Scatter plot showing the correlation of gene expressions between shCTCF#1 and shCTCF#2.

**Fig S2. Analysis of differentially expressed genes between WT cells and CTCF KD cells.**

(A) The expression of CTCF is correlated with the cell coordination on PC2.

(B) Differentially expressed genes were plotted in MAplot. Significantly up-regulated genes and down-regulated genes were indicated by red and blue, respectively.

(C) The top 15 enriched GO terms in the 195 up-regulated genes.

(D) The top 15 enriched GO terms in the 107 down-regulated genes.

**Fig. S3. Histogram showing that CV difference of gene expression between CTCF-KD cells and WT cells followed normal distribution.**

