## Supplementary figures and images for "Exploring the Changing Landscape of Cell-to-Cell Variation After CTCF Knockdown via Single Cell RNA-seq"

### Supplemental Figur1

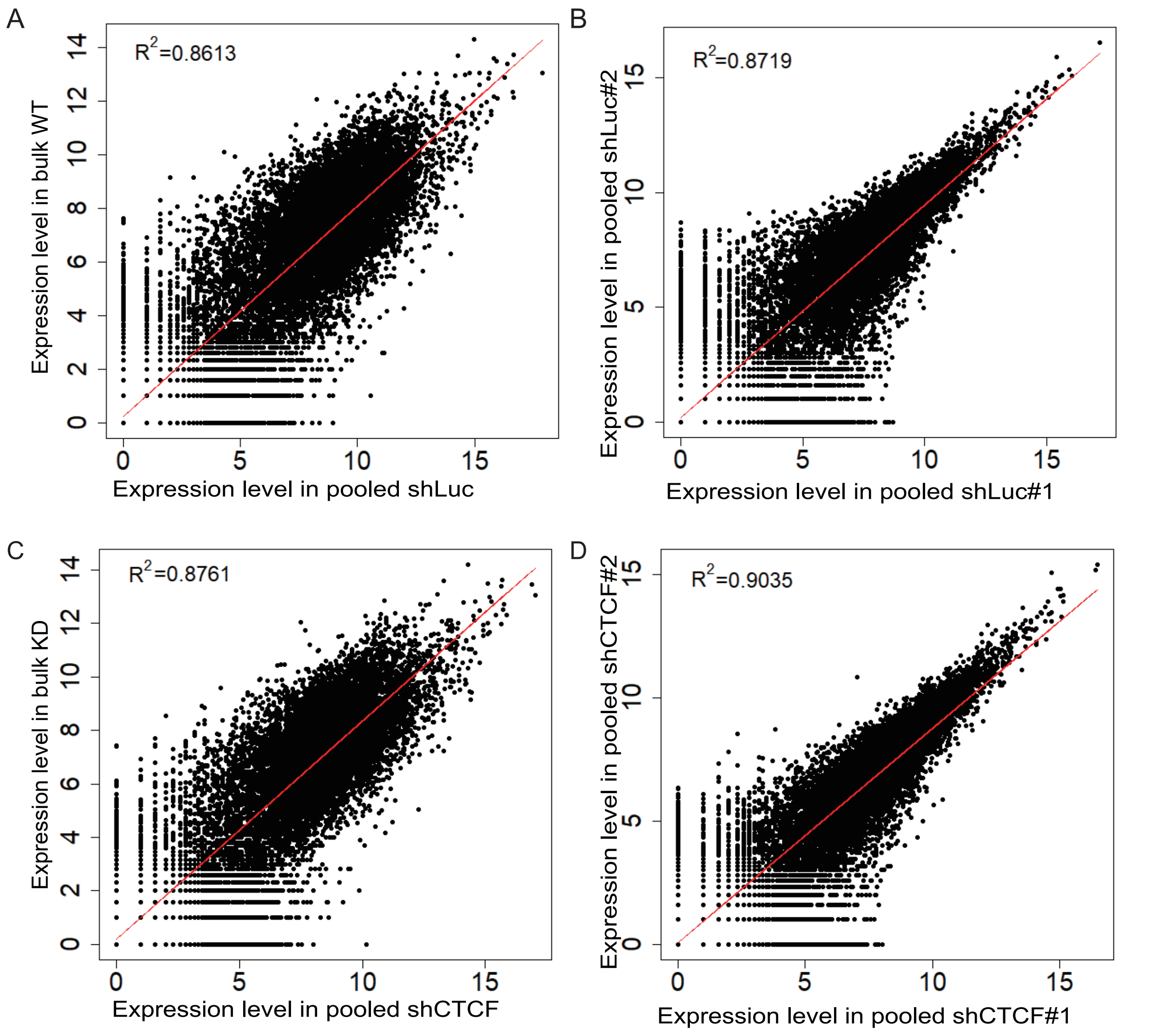

### Supplemental Figur3

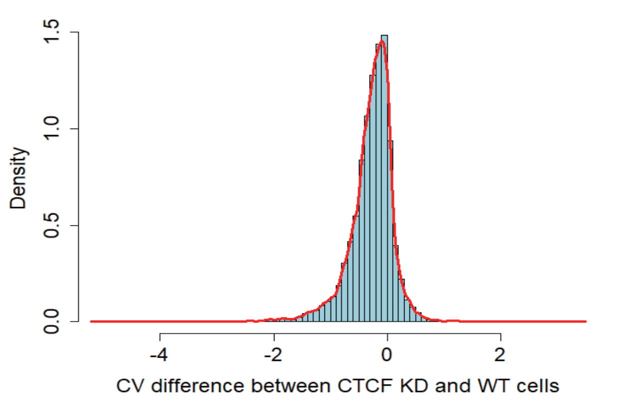

### Supplemental Figure2

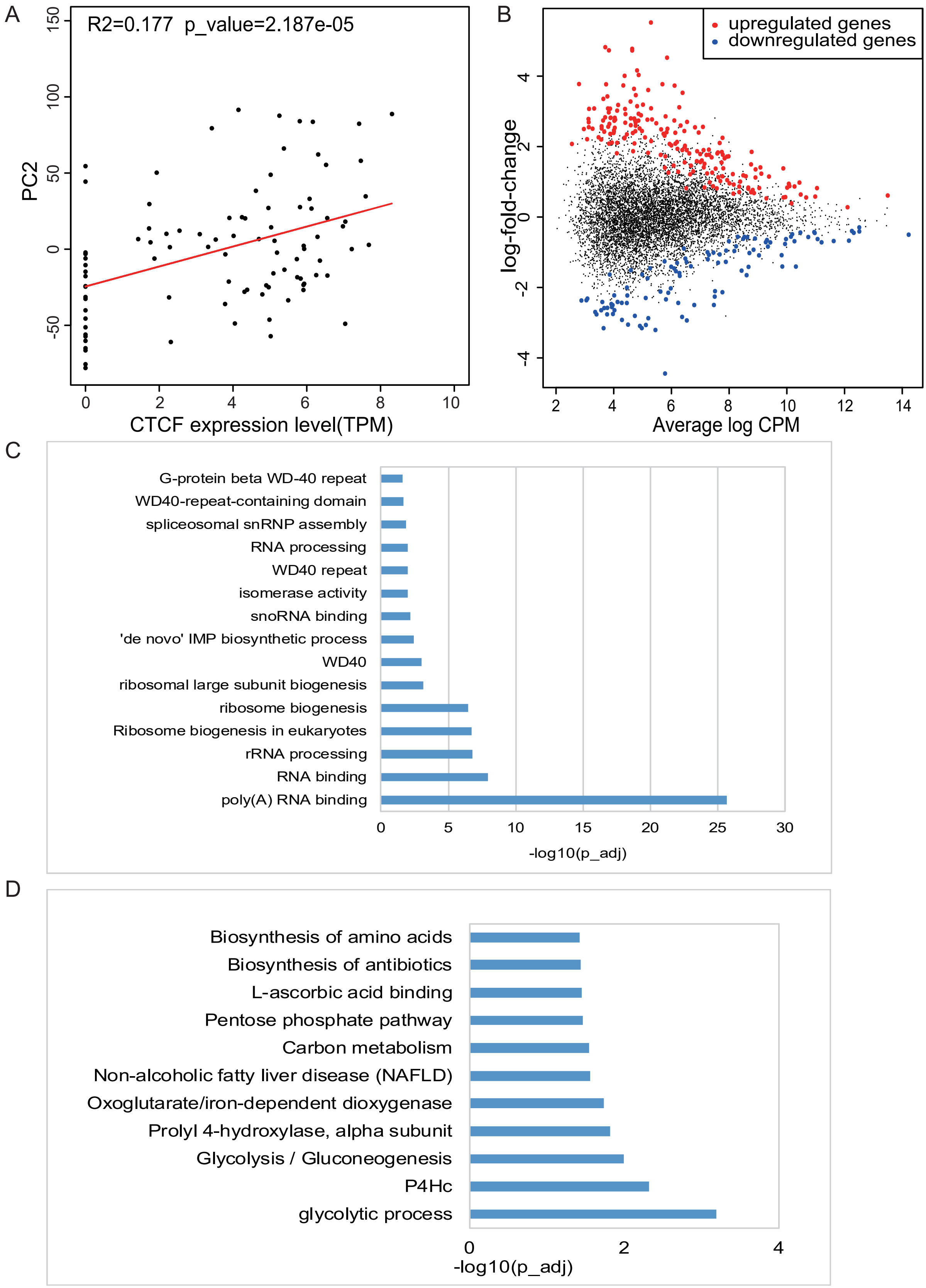
